# Somatic polyploidy supports biosynthesis and tissue function by increasing transcriptional output

**DOI:** 10.1101/2024.03.25.586714

**Authors:** Alexander T. Lessenger, Mathew P. Swaffer, Jan M. Skotheim, Jessica L. Feldman

**Affiliations:** Department of Biology, Stanford University, Stanford, CA 94305, USA; Wellcome Centre for Cell Biology, University of Edinburgh, Edinburgh, Scotland, EH9 3BF, United Kingdom; Chan-Zuckerberg Biohub, San Francisco, CA

**Keywords:** scaling, polyploidy, biosynthesis, intestine, *C. elegans*

## Abstract

Cell size and biosynthetic capacity generally increase with increased DNA content. Polyploidy has therefore been proposed to be an adaptive strategy to increase cell size in specialized tissues with high biosynthetic demands. However, if and how DNA concentration limits cellular biosynthesis *in vivo* is not well understood, and the impacts of polyploidy in non-disease states is not well studied. Here, we show that polyploidy in the *C. elegans* intestine is critical for cell growth and yolk biosynthesis, a central role of this organ. Artificially lowering the DNA/cytoplasm ratio by reducing polyploidization in the intestine gave rise to smaller cells with more dilute mRNA. Highly-expressed transcripts were more sensitive to this mRNA dilution, whereas lowly-expressed genes were partially compensated – in part by loading more RNA Polymerase II on the remaining genomes. DNA-dilute cells had normal total protein concentration, which we propose is achieved by increasing production of translational machinery at the expense of specialized, cell-type specific proteins.

## Introduction

Proliferating cells coordinate DNA replication with growth to maintain the ratio between the genome and cytoplasm volume. In the absence of DNA replication, the genome is diluted by cell growth, leading to biosynthetic collapse in extreme cases (Neurohr et al., 2019). Many organisms have adapted strategies to overcome this apparent limitation on cell size (Marshall et al., 2012; O’Farrell, 2015). The main such strategy is polyploidization, the process of generating more than two copies of the genome in a single cell. Most multicellular organisms have polyploid cell types, which arise during normal development by converting a mitotic cell cycle into an endocycle where DNA is replicated without subsequent cell division or by the fusion of many cells into a single syncytium (Edgar et al., 2014). These events are developmentally programmed in many tissues, such as the insect salivary gland, mammalian liver, and plant leaf epidermis (Dej and Spradling, 1999; Roeder et al., 2012; Gentric and Desdouets, 2014). Several studies have shown that increased DNA content causally increases cell size (Unhavaithaya and Orr-Weaver, 2012; Hansson et al., 2020; Ma et al., 2022; Darmasaputra et al., 2024). Given this association between DNA content and cell size, polyploidization may enable an accompanied increase in biosynthetic capacity, but such a requirement has never been tested *in vivo*.

A role for DNA content in facilitating rapid biosynthesis in multicellular systems would be consistent with work from proliferating, single-celled systems, where growth rate decreases as the genome dilutes (Neurohr et al., 2019). Proteins and transcripts change in concentration as the cell grows, but the proteomic and transcriptomic signatures of large cells are eliminated upon genome duplication (Neurohr et al., 2019; Chen et al., 2020; Lanz et al., 2022; Swaffer et al., 2023). This result, and the observation that polyploidization in cultured lymphocytic leukemia cells permits normal growth rate well beyond normal size (Mu et al., 2020), shows that these effects are due to the reduction of the DNA/cytoplasm ratio and not cell size *per se.* However, differentiated cells face substantially different pressures from single cells in culture that maximize their growth rate and divide as rapidly as possible, and thus whether these studies can be extrapolated to cells in multicellular organisms remains to be tested.

One major gap in our understanding is how DNA content mechanistically limits growth and biosynthesis. One possibility arises from recent work in budding yeast, showing that the concentrations of both RNA Polymerase II (Pol II) and the genome are limiting for mRNA transcription (Swaffer et al., 2023). However, only the few most highly expressed gene promoters may be near saturation, indicating that most genes have substantial capacity to load more polymerase and increase transcription (Swaffer et al., 2023). Thus, we hypothesize that only a few transcripts reach a gene copy number- and ploidy-imposed transcription limit, and that other transcripts scale their production accordingly.

The intestine of the nematode worm *Caenorhabditis elegans* offers an excellent model to test the importance of DNA content *in vivo* as it is a polyploid tissue with high biosynthetic capacity in a genetically tractable, multicellular organism. During normal *C. elegans* development, the intestine becomes polyploid through a series of programmed endocycles that are coordinated with progression through larval stages (Fig. 1 A; Hedgecock and White, 1985). In addition to its role in nutrient absorption and storage, the adult hermaphrodite intestine synthesizes an enormous amount of yolk protein, which is shuttled to oocytes and aids in the larval development of the next generation (Van Rompay et al., 2015; Perez and Lehner, 2019).

**Figure 1:**
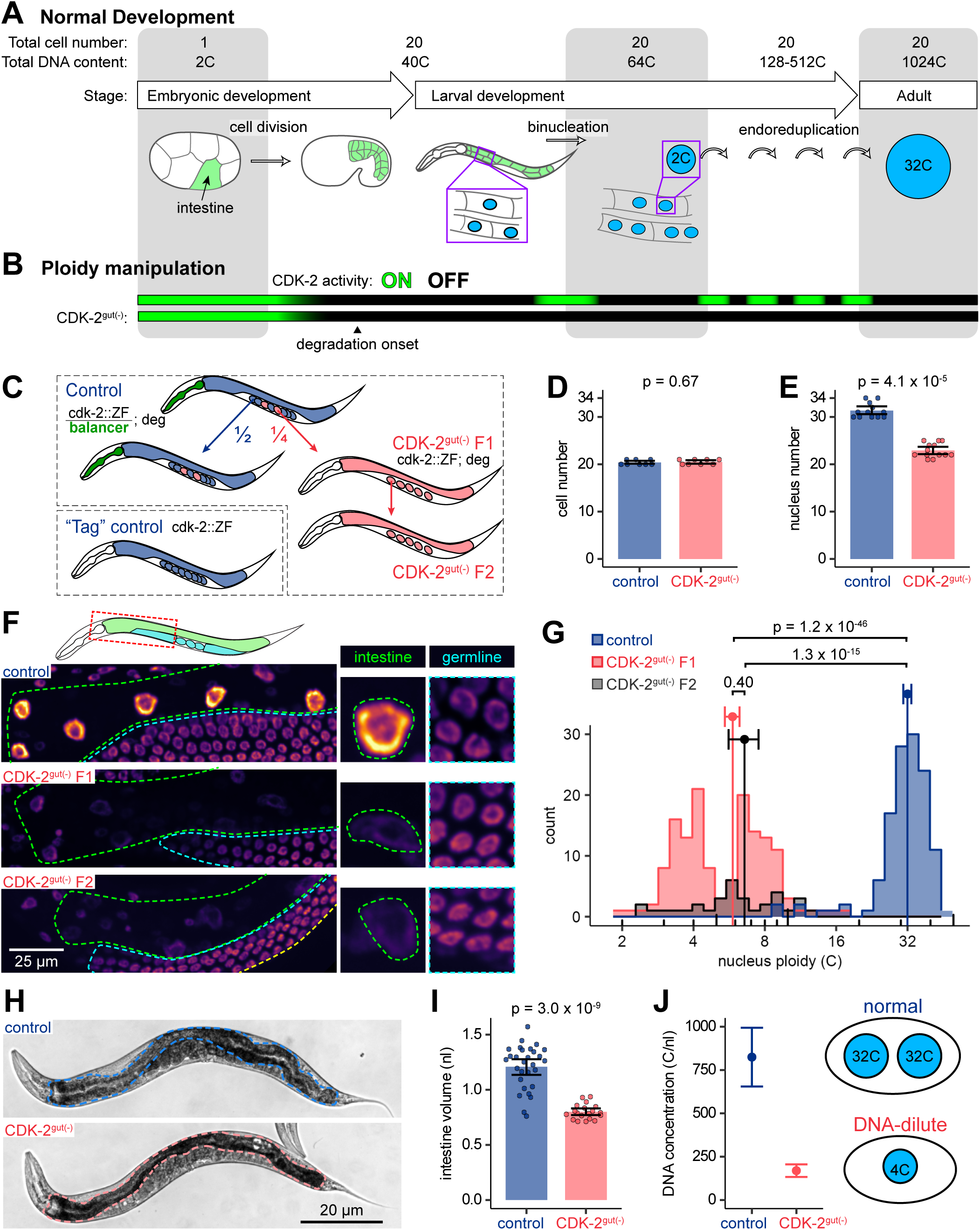
A tool to disrupt polyploidization. (A) Cartoon depicting *C. elegans* intestinal development, the origins of polyploidy in this tissue, and the cell numbers and ploidy values at each developmental stage. (B) Cartoon depicting timing of intes-tine-specific CDK-2 degradation and timing of CDK-2 activity in wild-type and CDK-2^gut(-)^ intestines. (C) Cartoon depicting genetics of experimental and control strains. (D) Quantification of intestinal nucleus number in L1 worms, representing final cell number. (E) Quantification of nucleus number in L4 worms, representing final nucleus number. (F) Sum projections of 3D images of day 1 adult worms fixed and stained with Hoechst DNA-binding dye (viridis-inferno coloring). Insets show nuclei from 3^rd^ intestinal ring and neighboring pachytene germline cells. Images are scaled so that germline nuclei appear at similar intensities because ploidy was quantified by normalized to these cells. Scale bar = 25 μm. (G) Quantification of nucleus ploidy in control and CDK-2^gut(-)^ F1 and F2 day 1-adult intestines. Dots and error bars represent means and 95% confidence interval. Kruskal-Wallis test p = 2.2 x 10^-48^; p values shown are from post-hoc Dunn test with Benjamini-Hochberg multiple hypothesis testing correction. (H) Representative DIC images of live adult control and CDK-2^gut(-)^ worms. Intestines are outlined in blue (control) or red (CDK-2^gut(-)^) dotted lines. Worms are age-matched at 96 hours post-egg lay. (I) Quantification of intestine volume, estimated from cross sectional area. Each point represents 1 worm. Volumes were estimated from length and cross-sectional area. In D, E, and H, bars and error bars represent mean and 95% confidence interval; P values from Mann Whitney U tests. (J) Left: estimation of DNA/cytoplasm ratio in control and CDK-2^gut(-)^ intestines from parameters in D-H. Error bars represent combined standard deviation. Right: cartoon depicting normal and DNA-dilute cells.

We successfully limited polyploidization of the intestine using tissue- and time-specific disruption of the cell cycle, allowing us to directly test the necessity of polyploidy for a variety of physiological functions, including cell and organelle size and the synthesis of yolk and other molecules. We demonstrate that polyploidy in the intestine is necessary for large cell size, but is not limiting for the final body size of the worm. Tissue-specific RNA-seq and Pol II ChIP-seq experiments revealed that reduced ploidy causes a global reduction in the expression of all genes, but has a more severe effect on the most highly expressed genes. Finally, we show that polyploidy is required for efficient biosynthesis of yolk protein and rapid development of progeny. Together, our data identify how biological molecules scale with a changing DNA/cytoplasm ratio in a multicellular, differentiated context, and demonstrate that polyploidy is a critical strategy for increasing biosynthetic capacity.

## Results and Discussion

### Intestine-specific depletion of CDK-2 disrupts polyploidization in the *C. elegans* intestine

We sought to develop a system where the ploidy of a polyploid tissue could be genetically limited. Polyploidy arises during the normal and stereotyped development of the *C. elegans* intestine. During embryogenesis, cell divisions increase intestinal cell number from 1 to 20 and total DNA content from 2C to 40C (C = 1 haploid genome equivalent). During larval development, most of these cells undergo endomitosis to become binucleated, followed by several rounds of endoreduplication that increase the ploidy of all individual nuclei from 2C to 32C, and of the whole organ to 1024C (Fig. 1 A; Hedgecock and White, 1985).

To modify intestinal ploidy, we degraded the G1-S promoting cyclin dependent kinase (CDK) CDK-2, which promotes entry into S phase in mitotic cell cycles and endocycles, using the ZIF-1/ZF protein depletion system (Armenti et al., 2014; Sallee et al., 2018). CDK-2 was endogenously tagged with a zinc finger degron (ZF), and a degrader (ZIF-1) was expressed in the intestine from the *asp-1p* promoter, which begins expression before binucleation but after the final intestinal cell divisions (Fig. 1 B). Worms with degraded CDK-2 are referred to as CDK-2^gut(-)^, and are the F1 homozygous progeny of balanced heterozygotes unless otherwise indicated (Fig. 1 C).

We found that CDK-2 degradation nearly eliminated endomitosis but did not affect cell division, decreasing the total nucleus number from 31.3 in control to 22.9, on average (Fig. 1 D and E). To determine whether CDK-2^gut(-)^ worms had reduced DNA content in intestinal nuclei, we quantified the total fluorescence intensity of nuclei after staining with Hoechst, relative to neighboring tetraploid pachytene germline nuclei. Average intestinal nucleus ploidy was reduced from 31.8C in control worms to 5.8C in CDK-2^gut(-)^ worms (Fig. 1, F and G). The bimodal distribution in CDK-2^gut(-)^ worms is partially explained by different ploidies along the anterior-posterior axis of the intestine, which may be due to different levels of ZIF-1 expression, or different sensitivities to the loss of CDK-2 due to their differing pattern of cell division/binucleation (Fig. S1 A). From the reduction in nucleus number and ploidy, we estimate that CDK-2^gut(-)^ intestines have a total DNA content of 134C, 13% of control.

Polyploidy has been shown to promote larger cell size in several contexts, so we estimated the size of the intestine from its cross sectional area and length to determine if the average cell size was limited in CDK-2^gut(-)^ intestines. We found that adult CDK-2^gut(-)^ intestines were 66% the size of control due to decreased width (Fig. 1, H and I and Fig. S1, B and C), despite having the same number of cells, showing that DNA content limits cell size in our system. It is important to note that because the reduction in DNA content is significantly larger than the reduction in cell size in CDK-2^gut(-)^ intestines, the DNA concentration in the intestine is diluted from 825 C/nl in control to 169 C/nl in CDK-2^gut(-)^ worms (Fig. 1 J). Thus, each copy of the genome supports a greater volume of cytoplasm, enabling cells to grow beyond their normal DNA/cytoplasm ratio and becomes DNA-dilute. Together, these data show that the intestine-specific depletion of CDK-2 severely limits DNA content and can therefore serve as a model to test the importance of high amounts of DNA in this tissue.

### Global increase in mRNA per genome buffers the effects of reduced ploidy

In budding yeast, DNA dilution ultimately causes cells to become diluted of RNA (Neurohr et al., 2019), and we wanted to see if this relationship also held in the *C. elegans* intestine. We measured global mRNA concentration by performing fluorescence in situ hybridization against the poly-A tail of mRNA (poly(A)-FISH), and found the cellular concentration of mRNA was reduced in CDK-2^gut(-)^ intestines relative to the neighboring germline (Fig. 2, A and B). However, this reduction in mRNA concentration (76% of control) was far less than the reduction in DNA concentration (21% of control), showing that there are active compensation mechanisms that increase the number of transcripts produced by each genome. One such mechanism identified in budding yeast is an increase in RNA Polymerase II (Pol II) abundance relative to the genome, which explains how transcription rates increase in larger yeast cells (Swaffer et al., 2023). To determine whether Pol II abundance similarly increased in DNA-dilute intestinal cells, we measured the amount of endogenously tagged Pol II (AMA-1::GFP) in the intestines of live worms. While nucleus size and total Pol II amount typically scale with cell size (Cantwell and Nurse, 2019; Swaffer et al., 2023), we found that nucleus size and Pol II abundance were maintained despite decreased cell size in CDK-2^gut(-)^ intestines (Fig. 2, D-G). The maintenance of Pol II levels in these DNA-dilute cells increased the Pol II-to-DNA ratio by 7.4-fold, which is expected to increase transcription from each genome. The transcription efficiency of Pol II could be further increased by the observed decrease in nuclear mRNA (Fig. 2 C), as nuclear mRNA inhibits Pol II transcription in cell culture (Berry et al., 2022).

**Figure 2:**
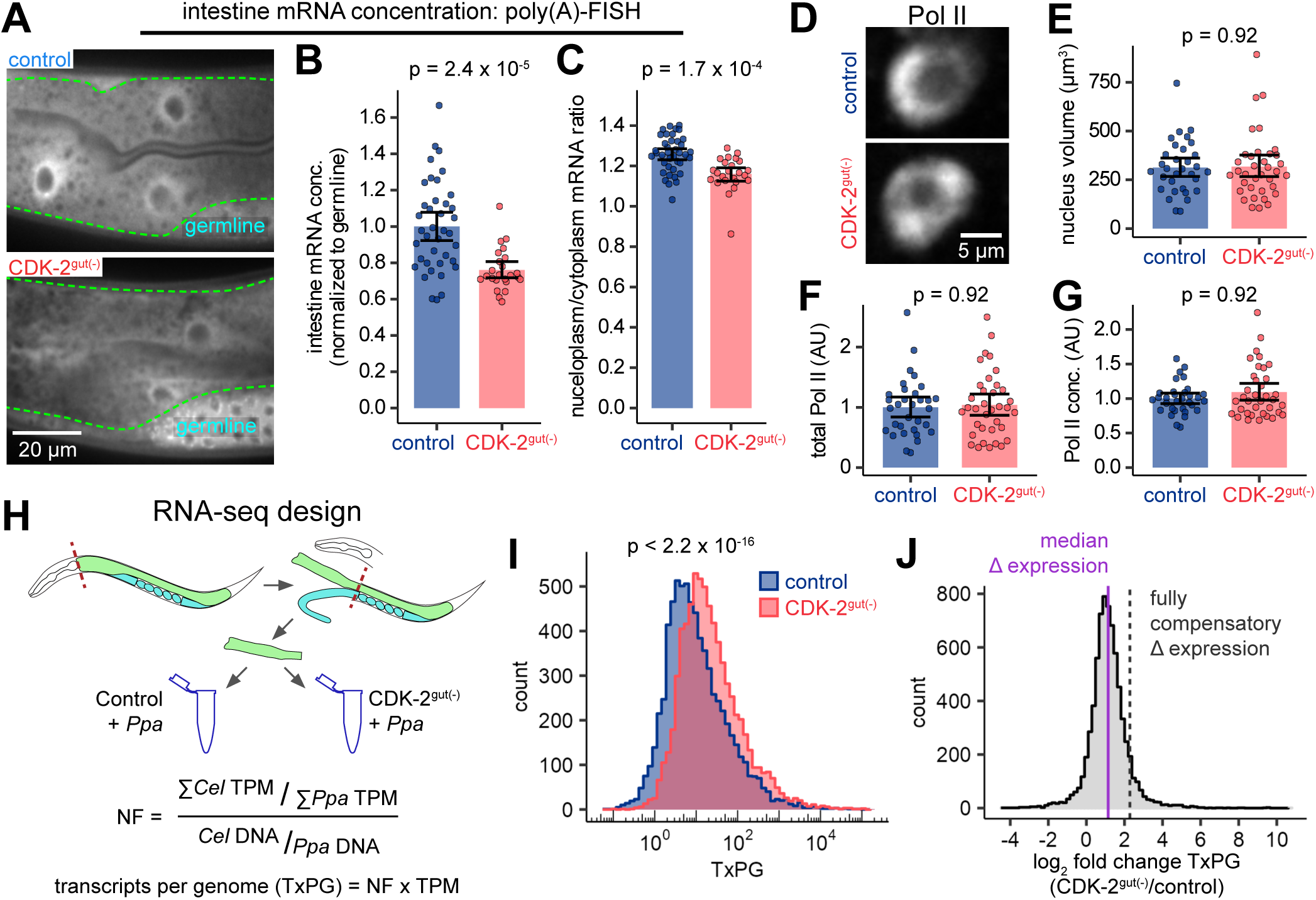
Scaling of cells and subcellular components with DNA content. (A) Confocal images of FISH against poly(A) tails in int2-int3 intestine rings of control and CDK-2^gut(-)^ in-testines. (B) Quantification of cytoplasmic mRNA concentration from images as in A. (C) Quantification of the ratio of the nucleoplasmic mRNA to the cytoplasmic mRNA from images as in A. (D) Live confo-cal images of intestinal nuclei of day 1 adult worms expressing endogenously-tagged Pol II (AMA-1::G-FP). Scale bar = 5 μm. (E-G) Quantification of the volume (E), concentration (F, mean fluorescence intensity) and total amount (G, total fluorescence intensity) of AMA-1. Each dot represents one nucleus, and bars and error bars represent mean and 95% confidence interval. P values from Mann-Whitney U tests after Benjamini-Hochberg multiple hypothesis testing correction. (H) Cartoon depicting normalized RNA-seq experiment: dissected intestines from two species were mixed, and *C. elegans* gene expression was normalized to the spike-in species, *P. pacificus*. (I) Distribution of transcripts per genome (TxPG) across all genes in control and CDK-2^gut(-)^ intestines. P value from t-test on the log of TxPG. (J) Distribution of changes in TxPG of all genes. Solid purple line represents the median change (Δ) in expression, and dashed black line indicates the level of expression change that would be necessary to fully restore normal mRNA concentration (1/ploidy reduction).

### Highly expressed transcripts are more sensitive to DNA limitation

Based on our observation that mRNA production globally increases in DNA dilute cells, we next sought to understand how DNA-dilution affects expression of individual genes, so we conducted a modified RNA sequencing (RNA-seq) experiment that could detect shifts in absolute RNA abundance. In standard RNA-seq methods, the measured abundance of a transcript is only meaningful in relation to other transcripts in the same sample, so we added a biological ‘spike-in’ which enabled us to directly compare absolute transcript concentration between samples. To each sample of dissected *C. elegans* intestines, we added equal amounts of dissected intestines of a related nematode, *Pristionchus pacificus* as a spike-in. *P. pacificus* was selected as a spike-in because it is expected to behave similarly to the *C. elegans* tissue at each step of sample processing. For each sample, we first calculated the global ratio of *C. elegans* to *P. pacificus* transcripts and then divided this ratio by the *C. elegans*-to-*P. pacificus* DNA ratio determined by qPCR. This normalization factor gives the relative ratio of total mRNA per genome in each sample, which can then be multiplied by the gene-specific quantification for each transcript (transcripts per million, TPM) to give a relative expression of the total number of transcripts per genome (Fig. 2 H). As we found that the DNA/cytoplasm ratio of CDK-2^gut(-)^ intestines was 21% of control (Fig. 1 J), these DNA dilute intestinal cells would need genes to increase transcripts per genome by 5-fold to fully compensate for the reduction in DNA concentration. However, we found that genes in CDK-2^gut(-)^ intestines had a median increase in transcripts per genome of 2.14-fold (Fig. 2, I and J), thereby corroborating our finding by poly(A)-FISH that DNA dilute cells increase RNA production, although not to a level that can fully compensate for the degree of DNA dilution.

How could the productivity of each genome increase? One possible model is that all genes compensate for limited DNA content by increasing transcription at a level inversely proportional to the level of DNA limitation (Fig. 3, A and B, ‘perfect compensation’ model, light red). Alternatively, the degree of compensation may depend on the initial expression levels (‘expression-dependent’ model, dark red) if highly expressed genes are in a regime where their promoters are closer to a saturation kinetic due to the initially high occupancy of Pol II slowing the loading of additional Pol II (Swaffer et al., 2023). This limitation would cause more highly expressed transcripts to decrease in abundance relative to control (‘expression-dependent’ model, dark red). Consistent with the expression-dependent compensation model, we found that the increase in transcripts per genome in CDK-2^gut(-)^ intestines decreased with increasing expression level (Fig. 3 C and Fig. S2 A), ruling out a model where limiting DNA template affects all transcripts equally. We considered the possibility that this effect may be caused by the activation of stress response genes, which are typically expressed at low levels, or by the altered expression of cell cycle-related genes. However, removal of either of these sets of genes from our analysis did not change the relationship (Fig. S2 B). Notably, our results do not imply a strict threshold for DNA limitation, below which increasing transcription perfectly compensates for limited DNA template. Instead, we observed a continuous decrease in compensation from lowly expressed to highly expressed genes.

**Figure 3:**
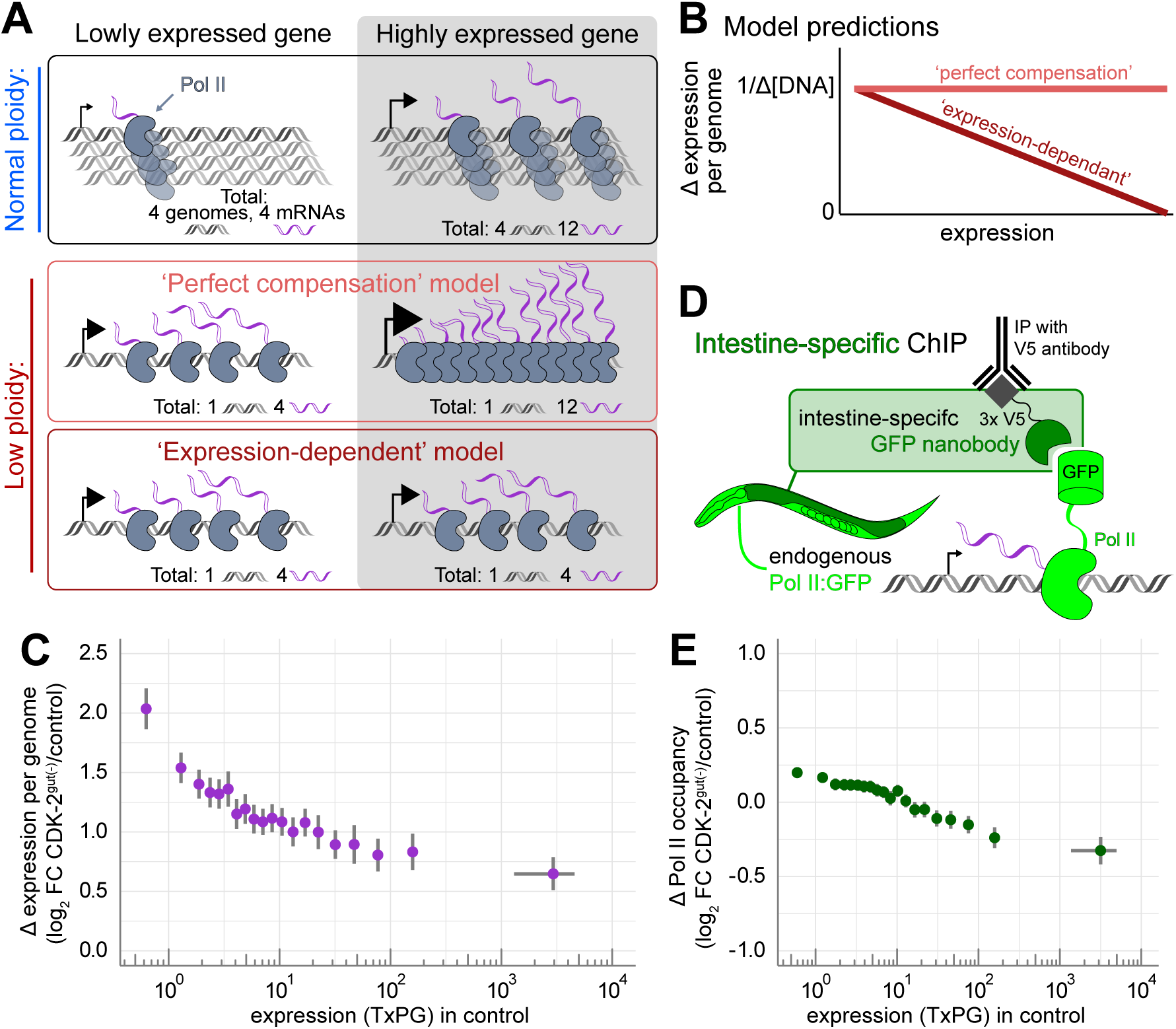
Highly expressed mRNAs are more limited by DNA content. (A) Cartoon depicting two models of transcriptional compensation for limited DNA content. (B) Mock data depicting how each model in A predicts how transcript abundance relative to genome copy number would change with increasing transcript abundance. (C) Change in spike-in normalized gene expression (TxPG) plotted against and binned by expression level in control, each bin represents 5% of genes. (D) Cartoon depicting experimental design for intestine-specific ChIP of Pol II. (E) Change in Pol II occupancy plotted against and binned by expression level in control, each bin represents 5% of genes. Dots in F and H represent mean fold change within the bin and error bars represent a 99% confidence interval.

### Differential impact of ploidy on highly and lowly expressed genes is partially mediated by RNA Polymerase II recruitment

We next wanted to determine whether the surprisingly high mRNA abundance in CDK-2^gut(-)^ intestines was achieved by increasing transcription rates or mRNA stability. RNA-seq is a measurement of RNA abundance, and so cannot distinguish between these models. To estimate the contribution of transcription rates, we performed a chromatin immunoprecipitation (ChIP)-seq experiment to quantify the amount of Pol II ‘occupancy’ on gene bodies specifically in the intestine. We expressed a GFP nanobody in the adult intestine from the *vit-5p* promoter in worms with endogenously tagged Pol II/AMA-1::GFP (Fig. 3 D). After cross-linking, we pulled down the nanobody to enrich for complexes of DNA, Pol II::GFP, and nanobody, which should only form in intestinal cells. Consistent with a transcription-dependent model, we observed a higher increase in Pol II occupancy on the gene bodies of more lowly expressed genes (Fig. 3 E and Fig. S2 C). However, the magnitude of the measured increase was less than that of the change in transcript abundance, suggesting that increased mRNA stability may also contribute to this compensation. It is also formally possible that linear changes in Pol II occupancy are not precisely quantified with our ChIP-seq method, leading to an underestimation of the degree of compensation due to transcription.

### Protein concentrations are buffered in low ploidy cells despite reduced mRNA concentrations

Together, our data support a model where several mechanisms work to buffer mRNA concentrations against loss of DNA template in low ploidy cells. However, this buffering compensation is not perfect as mRNA concentrations do decline in CKD-2^gut(-)^ intestines, just less than the template DNA concentration. Given this dilution of mRNA, we wanted to identify whether global protein concentration also decreased in this context, or whether further compensation mechanisms buffer protein levels. To estimate total protein concentration in the worm intestine, we stained fixed worms with a protein-binding dye (NHS-ester). Surprisingly, we found that the concentration of protein in the cytoplasm was unchanged between control and CKD-2^gut(-)^ intestines, relative to the neighboring germline (Fig. 4, A and B).

**Figure 4:**
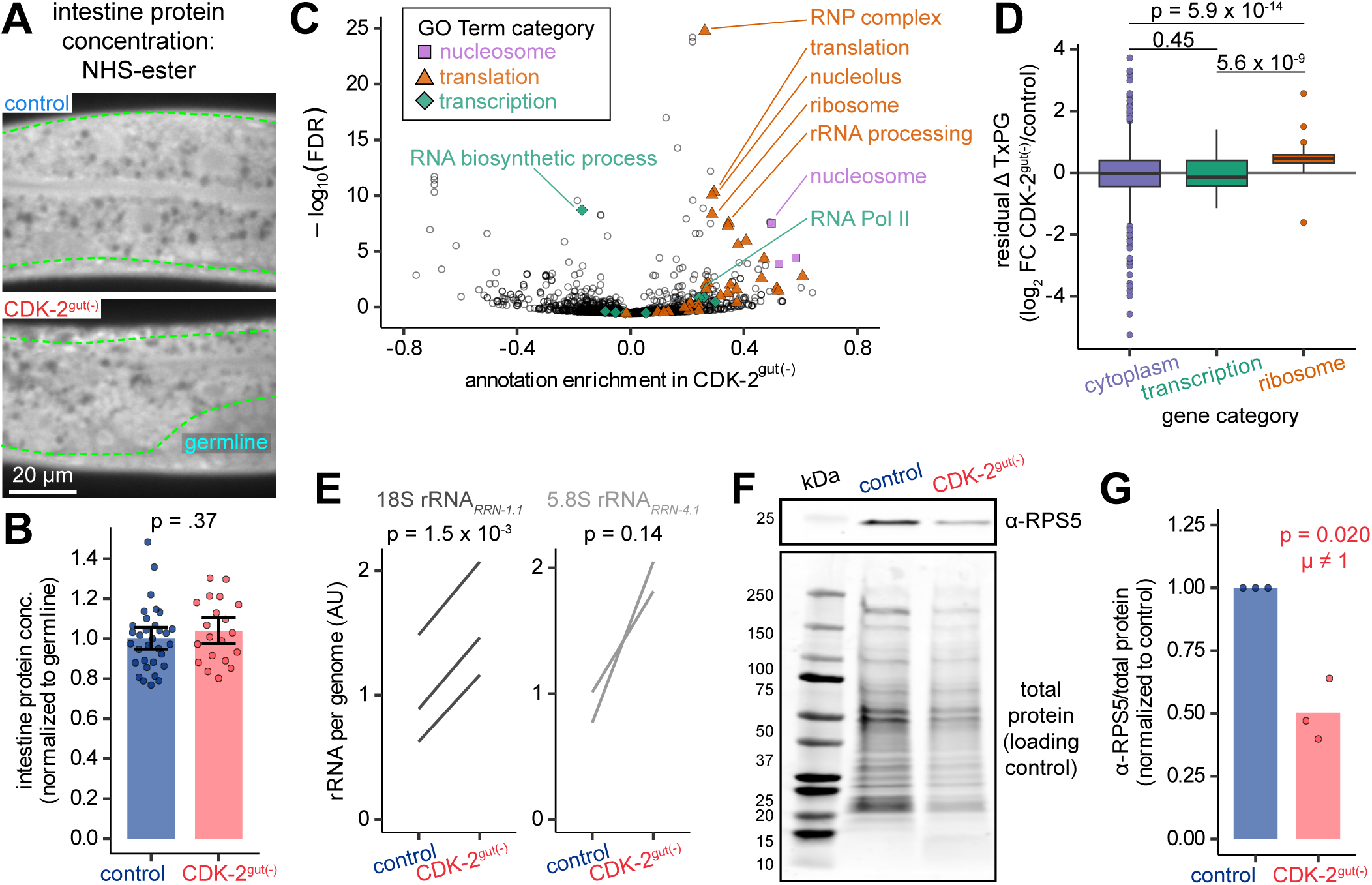
Ribosome production increases when DNA content is limited. (A) Confocal images of NHS-ester stain representing total protein in int2-int3 intestine rings of control and CDK-2^gut(-)^ intestines. (B) Quantification of cytoplasmic protein concentration from images as in A. (C) Volcano plot of GO terms for enrichment scores of transcripts in CDK-2^gut(-)^ intestines compared to control. Colored GO terms are related to the nucleosome (lavender squares), translation (orange circles), or transcription (green diamonds) (D) Box plots of residual change in RNA abundance after accounting for expression level (TxPG in control, relationship visualized in Fig. 3F). Log fold change in TxPG was regressed against expression level using a cubic polynomial. Groups are genes located in the cytoplasm (purple), involved in transcription (RNA Polymerase II subunit or general transcription factor; green) or ribosome subunits (orange). Kruskal-Wallis test p = 1.0 x 10^-13^; p values shown are from post-hoc Dunn test with Benjamini-Hochberg multiple hypothesis testing correction. (E) Quantification of change in rRNA per genome. Each line represents the change in TxPG of indicated ribosomal rRNA between paired RNA and DNA extractions from control and CDK-2^gut(-)^ intestines. Data are normalized to the mean TxPG of control. P values are of paired t test with Benjamini-Hochberg multiple hypothesis testing correction. (F) Representative western blot showing total protein and anti-RPS-5 signal in dissected control and CDK-2^gut(-)^ intestines. (G) Quantification of anti-RPS-5 signal normalized to total protein as in F, with control ratios for each blot scaled to 1. Each dot represents one experiment (N = 3 blots, n = 50-80 worms per genotype per blot). P value is of 1 sample t test against null hypothesis that μ = 1.

To understand how protein concentration can be kept constant when global mRNA concentrations are reduced, we investigated whether genes encoding translational machinery were up-regulated in our RNA-seq dataset. As an unbiased approach, we calculated an enrichment score for each gene ontology (GO) annotation that identifies annotations whose member genes are, as a group, more or less abundant compared to the global distribution (Cox and Mann, 2012). Consistent with the hypothesis that CDK-2^gut(-)^ intestines compensate for limited mRNA concentration by increasing translation rates, we found genes involved in translation to be significantly up-regulated (Fig. 4 C, orange). Despite the increase in mRNA per genome, we found terms related to transcription (green) were not uniformly up-regulated, consistent with a regime in which DNA content is much more limiting for transcription than Pol II. From this analysis we also found a strong up-regulation of nucleosome genes (lavender), which was surprising because histones, their main protein component, typically scale with DNA content (Claude et al., 2021; Swaffer et al., 2021). The divergence in the behavior of transcripts encoding transcriptional versus translational machinery was even more clearly seen after accounting for their relative expression levels: we regressed the fold change in transcripts per genome to the log of the expression level with a polynomial regression (R^2^ = 0.091, p = 2.2 x 10^-16^) and found that the residual change of ribosome subunits was significantly increased relative to those of Pol II and general transcription factors, or of all cytoplasmic genes (Fig. 4 D).

An increase in ribosome levels could further compensate for reduced mRNA by increasing translation, so we measured the volume of the nucleolus and the abundance of the pre-rRNA processing protein, Fibrillarin/FIB-1, both of which have been shown to correlate with ribosome production (Frank and Roth, 1998; Lee et al., 2012; Yi et al., 2015; Uppaluri et al., 2016). We found that the total amount of FIB-1 per nucleus was unchanged in CDK-2^gut(-)^ intestines, but its local concentration increased due to these cells having smaller nucleoli (Fig. S3).

To estimate the ribosome abundance more directly, we measured the change in ribosomal RNA (rRNA) and ribosomal protein subunit (RPS) RPS-5. Genome-normalized rRNA levels, as measured by qPCR, were increased in CDK-2^gut(-)^ intestines relative to control (Fig. 4 E). These measurements, combined with our estimate of DNA concentration, predict that ribosome concentration in CDK-2^gut(-)^ intestines should be 38% of control, despite the 5-fold reduction in DNA concentration. RPS-5 levels were measured in dissected intestines of control and CDK-2^gut(-)^ worms by western blot. Normalizing RPS-5 intensity to the total protein in the sample revealed that the fraction of the proteome dedicated to RPS-5 was decreased in CDK-2^gut(-)^ intestines to 50% of control (Fig. 4, F and G). The absolute reduction in rRNA and RPS-5 abundance in CDK-2^gut(-)^ intestines is reflective of a DNA-imposed limitation, but, like global mRNA, these ribosomal components are increased relative to DNA content.

Together, these data identify an additional layer of compensation where translational machinery is up-regulated relative to other genes of a similar expression level. However, this up-regulation can only partially explain how global protein concentration is maintained in DNA-dilute cells because ribosome concentration does decrease. Recent work *in vitro* has shown that protein concentration can be maintained by increasing translation efficiency relative to protein degradation as cytoplasm becomes more dilute (Chen et al., 2023). Thus, we speculate that increased translation efficiency and decreased protein degradation rates could also play a role in buffering protein concentration in DNA-dilute cells. However, because cell size is decreased and protein concentration is constant, the total protein amount is lower and therefore limited by DNA content.

### DNA content is required for a major physiological role of the *C. elegans* intestine

In a highly biosynthetic organ like the intestine, we expect protein synthesis to be bottlenecked at the ribosome, and so the increase in production of some proteins in CDK-2^gut(-)^ intestines, such as ribosomes themselves, may come at the expense of the production of other proteins. We therefore asked whether intestinal function was compromised in CDK-2^gut(-)^ worms.

To test whether the DNA-dilute intestine could meet the nutritional demands of the worm, we measured the maximum body size and growth rate. We found that maximum body size was only slightly reduced in CDK-2^gut(-)^ worms compared to control, and this difference was accounted for entirely by the reduced size of the intestine (Fig. 1 G and Fig. 5 A and B). The growth rate during development was also unaffected (Fig. 5 C), suggesting that DNA-dilute intestines are still able to meet the nutritional demands of organism growth during development and maintenance as an adult.

**Figure 5:**
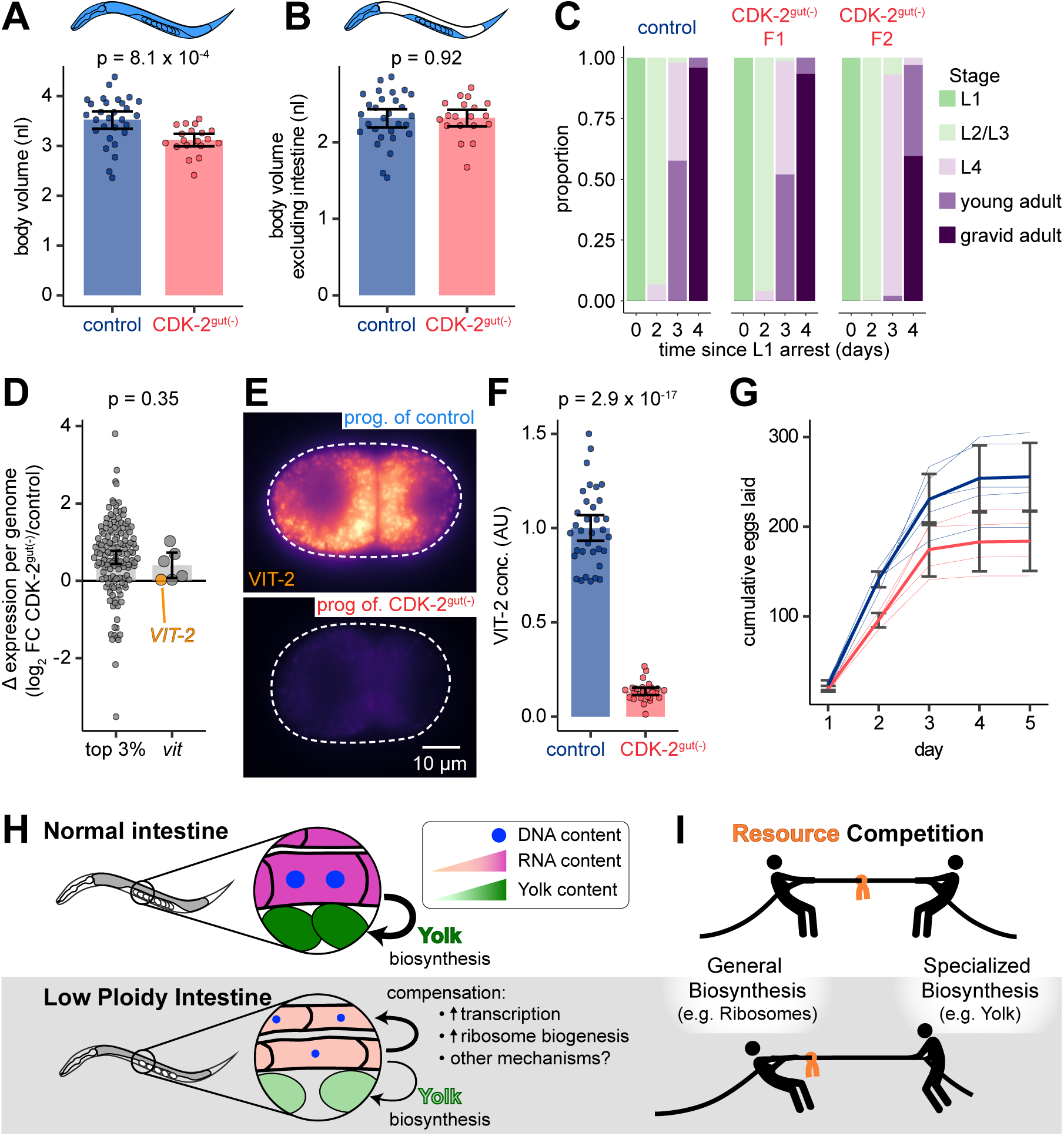
Polyploidy in the intestine is essential for biological fitness. (A and B) Quantification of body volume (A), and body volume excluding intestine (B) in control and CDK-2^gut(-)^ worms. Each dot represents 1 worm. Volumes were estimated from length and cross-section-al area. (C) Developmental progression rate of tag-control, CDK-2^gut(-)^ F1, and CDK-2^gut(-)^ F2 animals. Stacked bar graphs show the average proportion of worms at indicated developmental stage each day after release from overnight L1 arrest, grown at 15 °C (N=2 trials, n = 32-50 worms per genotype per trial). (D) Log_2_ fold change in TxPG of either the top 3% most highly expressed transcripts or the 6 vitel-logenin transcripts. Each dot represents 1 gene. (E) Representative widefield epifluorescence images of endogenously-tagged VIT-2::GFP in F1 progeny of control and CDK-2^gut(-)^ siblings. (F) Quantification of average VIT-2::GFP intensity in cytoplasm of 2 cell-stage embryos as in E. Each dot represents 1 embryo. In A, B, D, and F, bars and error bars represent means and 95% confidence intervals, and p-values are from Mann-Whitney U tests. (G) Cumulative number of eggs laid per day of adulthood in control and CDK-2^gut(-)^ animals. (H) Cartoon summarizing the cellular and organismal phenotypes of CDK-2^gut(-)^ worms (I) Cartoon depicting a hypothesized biosynthetic tradeoff between supporting the cell through general biosynthesis and supporting progeny through yolk production.

One of the main, and most biosynthetically demanding functions of the intestine is to produce vitellogenin yolk proteins (VIT) that are shuttled to the oocytes to aid the larval development of the next generation (Van Rompay et al., 2015; Perez and Lehner, 2019). Thus, we explored the effect of reduced polyploidy on the production of yolk protein and embryo. First, we examined the relative expression of transcripts encoding yolk proteins in our RNA-seq dataset. This analysis revealed that these transcripts are only slightly compensated, in line with other highly expressed genes (Fig. 5 D), causing their share of the transcriptome to decrease from 22% of total mRNAs to 15%. Unlike the compensation we saw for total protein, we found that CDK-2^gut(-)^ animals produce substantially less yolk protein than their control siblings (Fig. 5, E and F), which we measured in the early embryos of their progeny. Notably, this decrease is far more severe than the decrease in VIT-2::GFP in embryos of worms where the binucleation is converted to an endoreduplication (Rijnberk et al., 2022). These binucleation-defective animals also decrease expression of vitellogenin mRNA, but their progeny have ∼85% of control VIT-2::GFP signal. In contrast, embryos from CDK-2^gut(-)^ mothers have only 13% VIT-2::GFP compared to control, indicating that ploidy reduction limits protein synthesis far more severely than merely changing DNA compartmentalization.

Consistent with phenotypes of mutations that reduce yolk protein levels, CDK-2^gut(-)^ worms laid fewer eggs, and their progeny suffered developmental delays (Fig. 5, C and G). While CDK-2^gut(-)^ worms from control mothers (CDK-2^gut(-)^ F1s) developed at a normal rate, their progeny (CDK-2^gut(-)^ F2s) developed more slowly, suggesting that this slow growth phenotype stems from defects in yolk deposition into the embryos that gave rise to these worms. To rule out the possibility that the slower growth of CDK-2^gut(-)^ F2s was due to a difference in ploidy due to the inheritance of wild-type CDK-2 from the mothers of CDK-2^gut(-)^ F1s but not F2s, we measured ploidy in both of these genotypes. We observed no difference in ploidy between CDK-2^gut(-)^ F1s and F2s (Fig. 1 F), indicating that growth rate differences between F1s and F2s are due to differences in yolk content. These data indicate that ploidy reduction in the intestine severely impacts the specialized biosynthetic output of this tissue that supports progeny, constituting a severe fitness penalty to limiting DNA content in the intestine.

One of the main advantages of our multicellular model is that we can study the consequences of diluting DNA in a terminally differentiated cell, which is not optimized to divide as rapidly as possible, unlike the proliferating cells in which most DNA dilution models have been studied. We found that reduced polyploidy in the *C. elegans* intestine results in RNA dilution and a partially compensatory increase in production capacity of each genome, and that other, unidentified, molecular mechanisms stabilize protein concentrations (Fig. 5 H).

Importantly, these compensation mechanisms operate independently of the ‘sizer’ mechanism which couples cell size to DNA content in proliferating cells (Schmoller et al., 2015; Xie and Skotheim, 2020), and are thus compatible with terminally differentiated cells. However, we observed a 2-fold increase in mRNA and rRNA per genome, similar to what might be required to compensate for growth-mediated genome dilution over the course of a normal mitotic cell cycle. It is possible that yolk production is sacrificed in order to lessen the burden on translational machinery to maintain protein concentration and fulfill functions essential for organism viability (Fig. 5 I).

### Why would a tissue contain few big cells instead of many small cells?

Our results show that limiting DNA content restricts the biosynthetic capacity of the *C. elegans* intestine with profound physiological effects, suggesting that this tissue requires a minimum amount of DNA to properly function. Why might polyploidy have evolved in this context as a strategy to increase DNA content as opposed to increasing the number of diploid cells?

One explanation may be that nematode intestines face pressure to minimize cell number during embryonic and larval development. Previous work showed that increasing intestinal cell number during embryogenesis causes aberrant junction and apical surface formation and lumen deformation (Naturale et al., 2023; Demouchy et al., 2024). If unrepaired, extra junctions can cause lumen discontinuities, leading to starvation and arrest (Sallee et al., 2021). Efficient polarization of the embryonic intestine is thought to depend on the more stable contacts between pairs of intestinal cells than between intestinal cells and their non-intestinal neighbors (Naturale et al., 2023). This stability may be facilitated by the unusually large size and slow cell cycle of the intestine primordium. Together, these observations imply a maximum intestinal cell number for efficient embryonic organogenesis. So why doesn’t intestinal cell number increase after embryogenesis to meet the biosynthetic demands of the adult organ? Cell division has been shown to be an assault on tissue integrity once the intestine is polarized as mitosis breaks existing cell-cell junctions (Sallee et al., 2021). Mitosis in the larval intestine might be still more disruptive, since the structures are more mature, and such a disruption would make animals prone to infection as the intestine is actively passing food. Thus, larval intestinal divisions are likely highly disfavored as an evolutionary strategy to increase DNA content thereby requiring polyploidization to achieve this goal.

This selective pressure against too many cells can also be seen in other systems where cell number minimization may serve as a means to reduce the number of cell-cell contacts. For example, the mammalian placenta contacts the uterus with a single, giant syncytium. It is hypothesized that this continuous cell membrane helps resist invasion by maternal immune cells and pathogens that exploit cell junctions to breach epithelia (Zeldovich et al., 2013). In the *Arabidopsis* sepal, increased cell size due to polyploidy affects the curvature of the leaves, which is critical to protecting the developing bud (Roeder 2012). This curvature may be dependent on mechanical properties due to the amount of cell wall, which is minimized when cells are large. Therefore, many organisms have adapted somatic polyploid cells to increase cell size to solve a variety of challenges in cell coordination and tissue structure.

Together, our results show that somatic polyploidy facilitates the expression of highly abundant transcripts in contexts where additional cell division would be detrimental to an organism. More work is needed to fully elucidate the mechanism by which DNA is limiting and should investigate the temporal dynamics at play during compensation. Future studies should formally test whether multiple diploids can function as well as one polyploid cell, or whether such a strategy encounters additional challenges.

## Methods

### Experimental model details

*C. elegans* and *P. pacificus* were grown on standard NGM plates seeded with OP50 *E. coli* at 20 °C unless otherwise noted, according to standard laboratory practices (http://www.wormbook.org). The C terminus of the endogenous CDK-2 locus was edited using the SEC method using the guide sequence, 5’-ACCGCTGCTGAACAATCATC-3’ (Dickinson et al., 2015). Single copy transgenes were generated with the RMCE method (Nonet, 2020). All other strains were generated by genetic crosses.

### Microscopy equipment

Images used to measure ploidy, nucleus and nucleolus volume, and AMA-1 and FIB-1 abundance were acquired using a Zeiss Axio Observer Microscope (Carl Zeiss) with a PlanApoChromat 40x/1.3NA objective, Prime BSI Express sCMOS camera (Photometrics), and CSU-W1 spinning disk (Yokogawa), controlled using Slidebook software (Intelligent Imaging Innovations). Referred to as the ‘Zeiss spinning disk’ below.

Images used to measure mRNA and protein concentration were acquired using a Nikon Ti-E inverted microscope (Nikon Instruments) with a PlanApoChromat 40x/1.3NA objective, a Yokogawa CSU-X1 confocal spinning disk head, and an Andor Ixon Ultra back thinned EM-CCD camera (Oxford Instruments - Andor), controlled by NIS Elements (Nikon Instruments). Referred to as the ‘Nikon spinning disk’ below.

Images used to measure body and intestine volume and VIT-2 concentration were acquired using a Nikon Eclipse Ni upright microscope (Nikon Instruments) with a 10x/0.25NA 40x/1.3NA objective, Lumencore Sola Light Engine, and an Andor Zyla DG-152V-C1E-F1 5.5 Megapixel Front illuminated Scientific CMOS camera (Oxford Instruments – Andor), controlled by NIS Elements (Nikon Instruments). Referred to as the ‘Nikon widefield’ below.

### Counting intestinal nuclei

Fluorescent nuclei (histone::mCherry) were counted on our Nikon widefield microscope at the L1 stage to determine cell number, and at the L4 stage to determine final nucleus number. Live worms were imaged on 3% agarose pads and immobilized with 2 mM Levamisole.

### Ploidy quantification and estimation of total organ ploidy

Day 1 adult *C. elegans* were washed in M9 for 20 min to remove bacteria. Worms were permeabilized in PBS + 0.1% Tween 20 (PBST) + 0.1% Triton-X 100 for 10 min, washed 2x in PBST, and fixed overnight in freshly prepared Carnoy’s solution (60% ethanol, 30% chloroform, 10% glacial acetic acid) at 4 °C. Worms were rehydrated with 5 min washes at 4 °C in 70% ethanol, 30% ethanol in 1X PBS, and 1X PBS. Worms were stained with 500 ng/μl Hoescht 33342 for 1 hour at 37 °C in PBST, washed 3x for 10 min at room temperature in PBST, and mounted in VectaShield. Worms were imaged with our Zeiss spinning disk microscope with a 0.6 μm z-step. We excluded images of worms whose intestines were not oriented toward the coverslip.

Nuclei were segmented using StarDist (Weigert et al., 2020) software followed by manual correction. Crops of images of both low ploidy and control intestines were manually segmented using napari software to create ground truth training data. Images and labels were upscaled 4x in z to make nuclei more isometric, then a 3D StarDist model was trained using default parameters. StarDist predictions were downscaled 4x in z to generate labels with original dimensions. These labels were manually corrected, intestine and pachytene germline nuclei were identified, and the sum intensity of each nucleus was calculated using a custom python script with napari and napari-simpleitk-image-processing packages. Cytoplasmic background was obtained manually from unstained images. C values (ploidy) were obtained using a custom R script: cytoplasmic background was subtracted from nucleus intensity values, and then corrected intensity values of intestinal nuclei were normalized to those of tetraploid pachytene germline nuclei (C=4).

Total intestine ploidy was calculated as average nucleus number × average nucleus ploidy (C_nuc_). Intestine DNA concentration was calculated as total intestine ploidy (C_tot_) / average intestine volume (nl). The error of these measurements was combined as

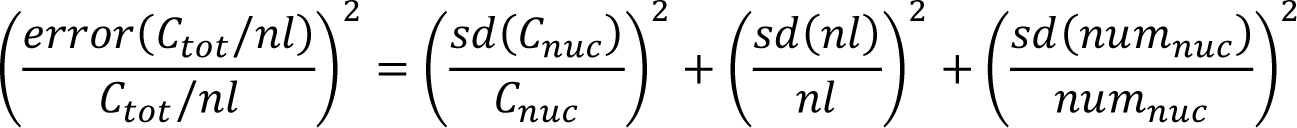

### Measurement of body and cell size

To measure maximum body and intestine size, day 2 adults (96 hours post egg lay) were imaged in bright field on our Nikon widefield microscope at 10X. Body and intestine were outlined in FIJI and the area and length were measured. Volume was estimated as a cylinder: V = π * (0.5*area/length)^2^ * length, since area is measured in the anterior-posterior x dorsal-ventral plane. Cell size was estimated as intestine volume/20.

### Measurement of AMA-1 and FIB-1 amount and volume (organelle size)

Worms expressing endogenously tagged AMA-1::GFP and wrmScarlet11::FIB-1 and an eft-3p::wrmScarlet1-10 transgene were imaged on our Zeiss spinning disk with a 0.3 μm z-step. Live worms were imaged on 3% agarose pads and immobilized with 2 mM Levamisole. Nuclei and nucleoli were segmented manually and their volumes and AMA-1 or FIB-1 intensities were calculated using the napari-simpleitk-image-processing package.

### Poly(A)-FISH and total protein image analysis and quantification

We discarded images of worms whose intestines were not oriented toward the coverslip. We manually segmented the nucleus and cell (and nucleolus for poly(A)-FISH) for each visible nucleus within the int2-int4 rings. Only the focal plane where each nucleus was largest was used for measurement. Several planes of the pachytene germline or proximal oocytes were also labeled for within-worm normalization. New segmentations were calculated for the cytoplasm (cell – nucleus) and nucleoplasm (nucleus – nucleolus). Background was estimated from unstained worms for poly(A)-FISH and from the slide for total protein. To calculate cytoplasmic intensity, background-subtracted intensities for each label were normalized to the germline signal in each worm. The mean of all cytoplasmic measurements was calculated for each worm and then scaled so that the mean of the control means was 1. To calculate the nucleoplasmic/cytoplasmic mRNA ratio, the nucleoplasm/cytoplasm ratio was calculated for each nucleus, then averaged for each worm, then scaled such that the mean of controls was 1. Images were segmented and measured using a custom python script with napari and simple-itk packages. 95% confidence intervals were calculated via bootstrap using the smean.cl.boot function in the R Hmisc package. Data were plotted in R.

### RNA-seq sample preparation

3 independent biological replicates were performed. DNA and RNA were extracted from dissected intestines. Day 1 adult (72 hrs post egg lay) *C. elegans* and *P. pacificus* (96 hrs post egg lay) were dissected and immediately flash frozen in liquid nitrogen. To dissect, worms were picked into drops of 340 mOsm L-15/FBS media (Fernandes Póvoa 2020) with 3 mM Levamisole to paralyze on a slide covered with Scotch tape. Working in batches, worms were cut behind the pharynx with a 25-gauge needle, then the intestine was cut away from the body. Intestines were washed 1x in L-15/FBS media to remove traces of other tissues, then transferred to PCR tubes and flash frozen in liquid nitrogen. Tissue was thawed directly into Trizol and lysates from each genotype were combined. An equal amount of *P. pacificus* lysate was distributed between tag control and CDK-2^gut(-)^ F1 lysates (N = 3 experiments, n ≈ 40 *P. pacificus* and either 80 control or 240 CDK-2^gut(-)^ F1 intestines per sample. Samples were split to separately extract RNA and DNA: RNA was extracted using Zymo Direct-zol Microprep kit, and DNA was extracted using Zymo DNA-quick microprep plus with a 4:1 DNA binding buffer-lysate ratio. RNA quality was assessed on a BioAnalyzer (RIN range from 8.7-10). RNA-seq libraries were prepared using NEBNext Single Cell/Low Input RNA Library Prep Kit for Illumina (NEB). Pooled libraries were sequenced with paired end sequencing on a Novoseq 6000 by Novogene.

### rRNA and DNA measurements

DNA and rRNA amounts were quantified by qPCR from the spike-in-normalized DNA and RNA samples described above. For rRNA quantification, cDNA was synthesized using the Superscript IV kit and random hexamers. qPCR was performed in technical triplicate using the PowerTrack SYBR Master Mix kit with primers against the *C. elegans* 18S and 5.8S rRNAs (*rrn-1.1* and *rrn-4.1,* respectively) and the *P. pacificus* 18S rRNA (*Ppa-rrn-1.1*). qPCR was performed similarly for DNA quantification using primers against a repetitive region of the *C. elegans* genome and the rDNA (*Ppa-rrn-1.1)* of *P. pacificus.* RNase and DNase treatments of DNA and RNA samples, respectively, ensured that rRNA and rDNA did not erroneously contribute to quantification of the other. Standard curves were made using these primers using DNA and RNA template prepared similarly to above and confirmed that these primers uniquely amplified targets from each species. For each sample, a DNA normalization factor was calculated from the DNA Cq values as 2^−𝛥𝐶𝑞(𝐶𝑒𝑙−𝑃𝑝𝑎)^, which reflects the ratio of *C. elegans* to *P. pacificus* DNA. For each rRNA, the rRNA per genome was calculated from the RNA Cq values and DNA normalization factor as 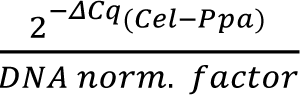. A paired t-test was performed for each rRNA experiment and corrected p-values with a Benjamini-Hochberg multiple hypothesis testing penalty.

### RNA-seq analysis

Fragments were aligned to concatenated WS284 *C. elegans* and *P. pacificus* genomes using STAR aligner with default parameters. Aligned reads were counted for each gene with featureCounts in R without multimappers. Counts were converted to transcripts per million (TPM) with the countToFPKM package in R. Transcripts per genome (TxPG) for each gene was then calculated by multiplying the TPM by a sample normalization factor:

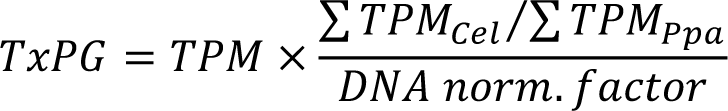

Within each genotype, we calculated the geometric mean of TxPG values for each gene, excluding genes with TxPG=0 in any sample, and then calculated the CDK-2^gut(-)^-to-CDK-2::ZF::GFP tag control ratio of the geometric means. We report the log_2_ of these fold changes. Genes were excluded from analysis if they had few reads across samples (DESeq2 baseMean<10) or were not expressed in the intestine. Genes were categorized as being expressed in the intestine if they had TxPG_control_>2 or were identified as expressed in the intestine by Kaletsky et al., 2018.

We quantified the trend in Fig. 3 F through a polynomial regression with 4 terms (y ∼ ax^0^ + bx^1^ + cx^2^ + dx^3^) in R. To test if this trend was caused by aberrant expression of stress response or cell cycle genes, we repeated that analysis after excluding genes with either GO biological process (GOBP) term “response to stress” or “cell cycle”. We further analyzed enrichment of GO terms using the 1D enrichment function in Perseus software (Cox and Mann 2012). We included all GO terms and keywords for the enrichment analysis and hypothesis testing, but limited our further analysis to GOBP and GO cellular compartment (GOCC) terms, which were the most interpretable.

The fraction of the transcriptome comprised of vitellogenin transcripts was calculated from the unadjusted TPMs in each sample as: ∑ 𝑇𝑃𝑀*_𝑉𝐼𝑇_*⁄∑ 𝑇𝑃𝑀*_𝑛𝑜𝑛−𝑉𝐼𝑇,𝑖𝑛𝑡−𝑒𝑥𝑝𝑟𝑒𝑠𝑠𝑒𝑑_*. The geometric mean of this fraction is reported for each genotype.

### ChIP-seq sample preparation

4 control and 3 CDK-2^gut(-)^ biological replicates were performed. For each replicate, 450,000 (control) or 600,000 (CDK-2^gut(-)^) age synchronized adult worms were flash frozen in liquid nitrogen. To synchronize, we picked several L4, balancer-negative worms (P0, JLF1584 control or JLF1585 CDK-2^gut(-)^) to NGM plates. Worms were grown at 15 °C to aid in monitoring of growth rate. Populations were allowed to starve, then starved F2 L1s were plated onto new 15 cm NGM peptone-rich plates seeded with NA22 *E. coli* (XL plates) at 30,000 (control) or 45,000 (CDK-2^gut(-)^) worms per plate. Gravid adults were bleached until most worms burst open (6-7.5 minutes vortexing at room temperature in solution of 2% sodium hypochlorite + 300 mM KOH), washed 4x in M9, resuspended in egg buffer, filtered through a sterile, 40 μm cell strainer, and hatched overnight at room temperature. Newly synchronized F3 L1s were again plated on XL plates and allowed to grow until adulthood when they were washed in cold M9, resuspended in PBS with protease inhibitors (PI; 2X Halt Protease Inhibitor Cocktail + 2 mM PMSF), and dripped into liquid nitrogen to form worm ‘Dippin’-dots’. Frozen worms were ground with a mortar and pestle, cooled with liquid nitrogen.

Our ChIP protocol was based on the protocol by Sen et al., 2021, in which buffer compositions and recipes can be found. Worm powder was thawed directly into PBS + PI + 1.1% formaldehyde and cross-linked for 7 minutes at room temperature. Working on ice from this point, each sample was pelleted, resuspended in B-ChIP-L0 to a final volume of 1.6 ml, and divided between 4 screw cap tubes. 0.8 ml ceramic beads were added to each tube, and worm pieces were lysed with a Fast Prep (3 cycles of 30 s at 5.5 M/s). Lysate was combined into 2 microcentrifuge tubes, brought up to 1.2 ml volume with B-ChIP-L0, and sonicated with a Branson 250 Sonifier. Lysates were cleared by centrifuging 2x 5 min at 20,000 g. Affinity resin was prepared at room temperature: 100 μl Protein G Dynabeads (Invitrogen) were washed 3x in B-ChIP-BL (0.5 mg/ml BSA in PBS), blocked in B-ChIP-BL for 40 minutes, bound to 19 μg anti-V5 antibody (SV5-Pk1, Bio-Rad), and washed 3x with B-ChIP-BL. Affinity resin was incubated overnight at 4 °C with the cleared lysate, then washed 2x with low salt wash buffer, 1x with high salt wash buffer, 2x with LiCl-containing wash buffer, and 2x in TE + 50 mM NaCl. On a ThermoMixer (Eppendorf) at 800 rpm, chromatin was eluted in B-ChIP-EL for 15 minutes at 65 °C, then treated with 0.4 mg/ml RNase A for 1 hour at 37 °C followed by 0.4 mg/ml proteinase K for 2 hours at 55°C, and finally cross links were reversed overnight at 65 °C. DNA was cleaned using the ChIP DNA Clean & Concentrator (ZYMO). Libraries were prepared using NEBNext Ultra II DNA Library Prep Kit for Illumina (NEB). Libraries were pooled and sequenced with paired end sequencing on a Novoseq 6000 (Novogene).

### ChIP-seq analysis

Fragments were aligned to the WBPS18 *C. elegans* genome using bowtie2. We calculated coverage of the 50 bp mid-point of each uniquely mapped fragment (*i.e.*, midpoint ± 25 bp) over all gene bodies using custom-written Python scripts, calculating both counts and fragments per kilobase million (FPKM) for each gene. Samples were excluded if the distribution of FPKM values was not highly bimodal, which indicated poor enrichment of intestine-specific gene bodies during the pull down. The geometric mean of FPKM values was calculated for control samples. Differential occupancy was calculated from the count data using DESeq2. Genes were excluded from analysis if they had few reads across samples (DESeq2 baseMean<10) or were not highly occupied in the intestine. Genes were categorized as being expressed in the intestine if they were identified as expressed in the intestine by Kaletsky et al., 2018 or had either baseMean or control FPKM values greater than the average of genes not expressed in the intestine in our RNA-seq experiment. The fold change from DESeq2 was then plotted against either the expression level in control, calculated from our RNA-seq data, or the occupancy in control, calculated from FPKM values from our ChIP-seq data.

### Estimation of ribosome concentration via western blot

Intestines from day 2 adult worms were dissected into egg buffer and flash frozen in liquid nitrogen. Intestines were thawed into 1x Laemmli sample buffer (Bio-Rad), pooled (40 for control, 80 for CDK-2^gut(-)^), and incubated 10 min at 95 °C. Lysates were loaded onto a 4-15% SDS polyacrylamide gel (Bio-Rad) and run for 1 hour at 120 V. Protein was transferred to a nitrocellulose membrane overnight in Towbin’s transfer buffer at 25 V on ice. Membranes were stained with REVERT total protein stain (LI-COR) and imaged in the 700 nm channel on an Odyssey DLx Imager (LI-COR). Membranes were blocked in 5% non-fat, dry milk in PBST, then stained with mouse 1:200 anti-RPS5-Alexa Fluor 546 (1°; sc-390935-A546, Santa Cruz Biotechnology; Nousch, 2020) followed by 1:10,000 IRDye 800CW goat anti-mouse IgG (2°, 926-32210, LI-COR). The A546 fluor was not used for quantification. Membranes were imaged again for the 800 nm channel. Images were processed in FIJI. Background was subtracted, and total RPS-5 signal was divided by total protein signal for each lane as a metric for the ribosome fraction of the proteome. Since there is, by definition, no variation in the control RPS-5/total protein, we performed a 1 sample t-test on CDK-2^gut(-)^ ratios against the null hypothesis that mean (μ)=1.

### Vitellogenin concentration

Age synchronized, gravid adults (72 hrs after egg lay) were picked into a 30 μl drop of M9 on a coverslip, then immobilized by placing the slide on a T-75 cell culture flask filled with ice water, then cut with a No. 10 scalpel to release eggs. Eggs were mixed 1:1 with 50% Iodixanol (OptiPrep, Sigma) to better match the refractive index of the egg (Xiong and Sugioka, 2020).

Coverslips were inverted onto slides and sealed with petroleum jelly. 1-4 cell-stage embryos were imaged with our Nikon widefield using a 60x/NA1.4 objective in the focal plane where the nuclei were largest.

### Brood size

L4 worms (48 hours post egg lay) were picked to individual, lightly seeded NGM plates (50 μl stationary phase OP50 grown for 6 hours). Each day after, adults were transferred to new plates and embryos were counted.

### Growth rate

Worms were grown at 15 °C. Gravid adults (P0) were bleached in 10 μl of bleach solution (2% sodium hypochlorite + 0.5 M KOH) on unseeded NGM plates. The next day, arrested F1 L1s were transferred to new, seeded plates. Every day after, we counted the number of worms in each developmental stage and transferred L4s to new plates to avoid confusing the F2 progeny of fast growing F1s with slow growing F1s.

## Acknowledgements

This work was supported by CMB Training Grant T32 GM007276 and a Stanford Graduate Fellowship awarded to A.T.L., an NIH New Innovator Award DP2 GM119136-01 and RO1 GM133950 awarded to J.L.F, and GM134858 awarded to J.M.S. Some strains were provided by CGC, funded by the NIH Office of Research Infrastructure Programs (P40 OD010440). We thank Georgi Marinov for sharing of code and Alex Long and Tim Stearns for use of their confocal microscope. We thank Lucy O’Brien, Lauren Cote, Andrew Fire, Hannah Fung, Kang Shen, Shicong Xie, and members of the Feldman and Skotheim labs for scientific discussions and feedback. We thank Dominique Bergman, Lauren Cote, Hannah Fung, Rachel Ng, Nabor Vasquez-Martinez, and Shicong Xie for their comments on the manuscript.

## Author Contributions

Conceptualization, A.T.L., J.L.F., and M.P.S.; methodology, A.T.L., M.P.S., and J.L.F.; software, A.T.L.; formal analysis, A.T.L.; investigation, A.T.L.; writing – original draft, A.T.L. and J.L.F.; writing – review & editing, A.T.L, J.L.F., M.P.S., and J.M.S.; visualization, A.T.L.; supervision, J.L.F. and J.M.S.; and funding acquisition A.T.L., J.L.F., and J.M.S.

---

**Figure S1:**
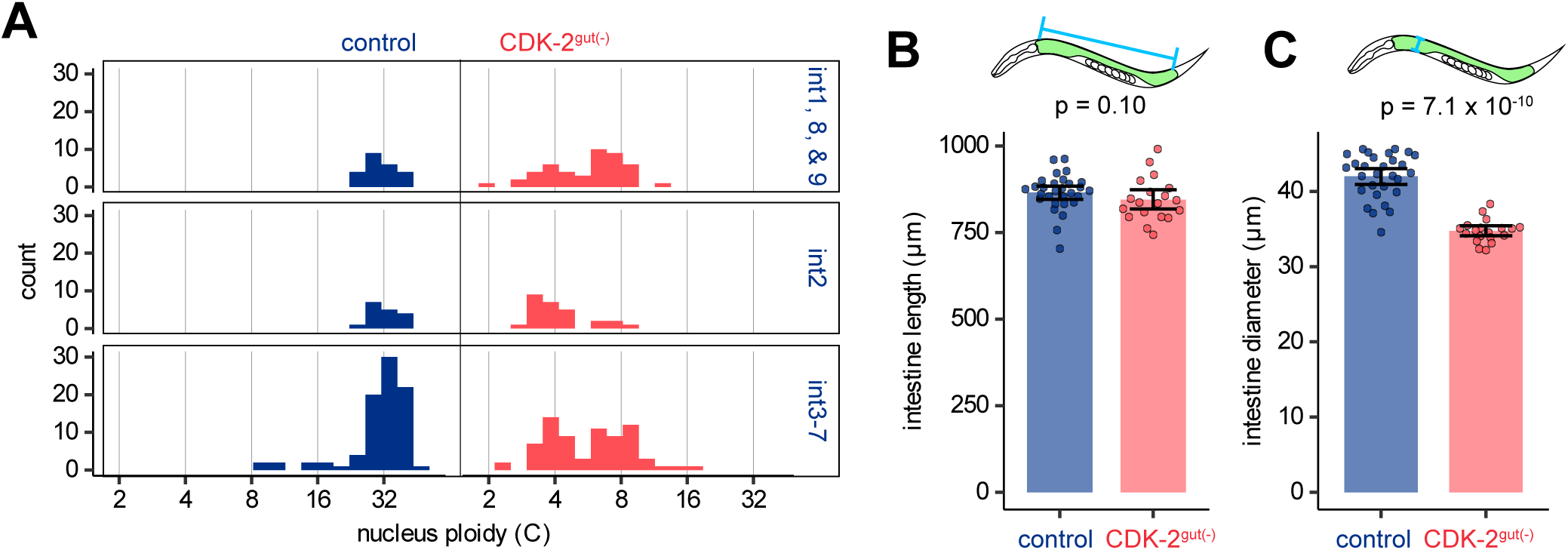
Extended data used in calculation of DNA/cytoplasm ratio. (A) Quantification of ploidy in control and CDK-2^gut(-)^ intestinal nuclei, grouped by intestinal subtype. (B and C) Quantification of intestine length (B) and width (C). In all graphs, each point represents 1 nucleus or worm, bars represent means and error bars represent 95% confidence intervals, and p-values are from Mann-Whitney U tests.

**Figure S2:**
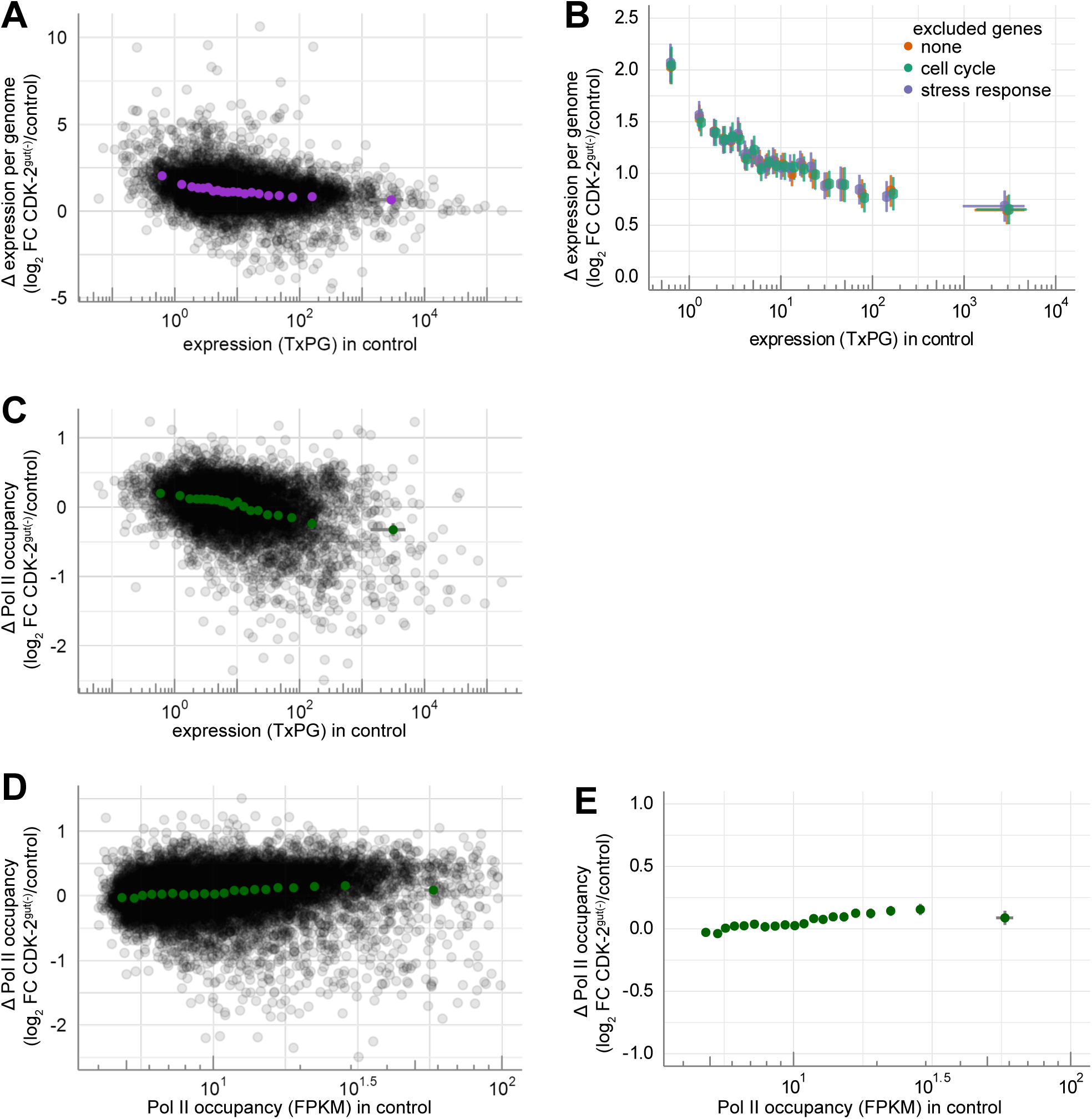
Extended presentation of RNA-seq and ChIP-seq data. (A) Presentation of full spread of individual genes from RNA-seq along with their binned averages from Fig. 3 F. (B) Change in spike-in normalized gene expression (TxPG) plotted against and binned by expression level in control, with and without removal of genes involved in stress response or cell cycle. Each bin represents 5% of genes. Genes were excluded before binning. (C) Presentation of full spread of individual genes from ChIP-seq along with their binned averages from Fig. 3H. (D and E) Change in Pol II occupancy plotted against and binned by occupancy in control, with (D) and without (E) individual genes. Each bin represents 5% of genes. In all graphs, black dots represent individual genes, colored dots represent mean fold change within a bin and error bars represent a 99% confidence interval.

**Figure S3:**
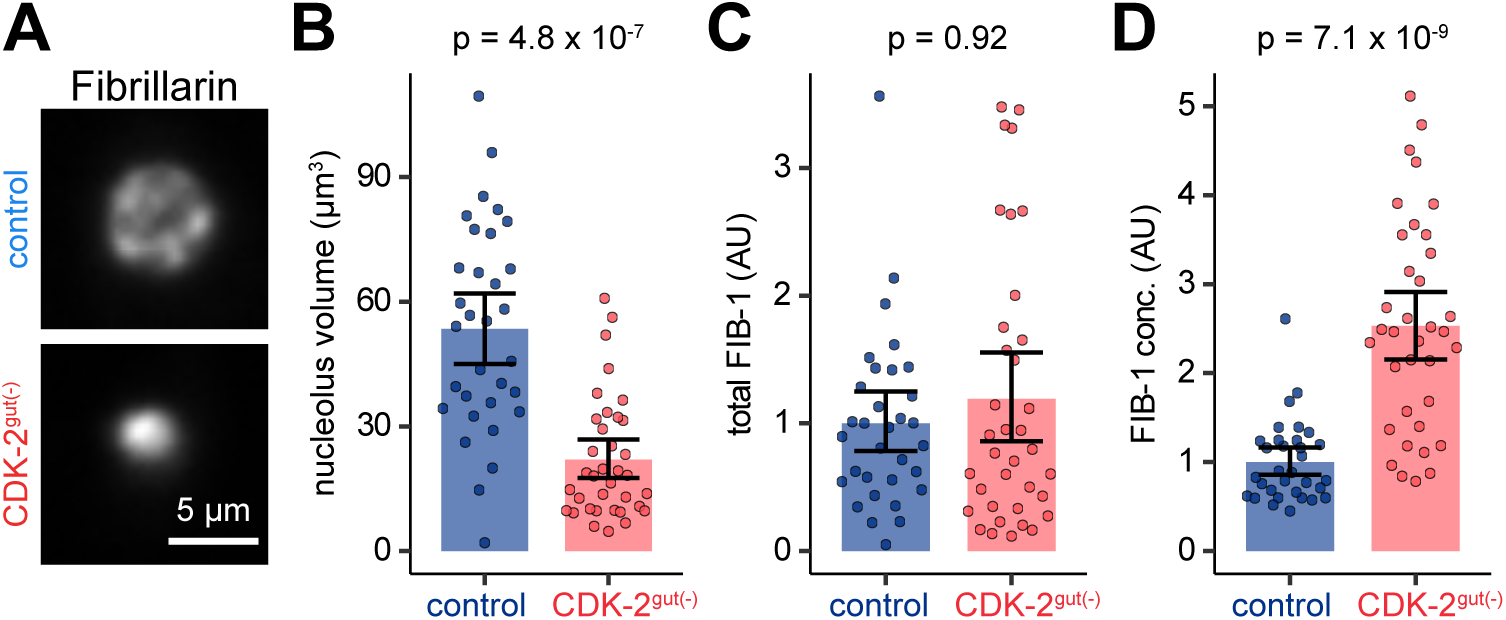
Quantification of nucleolus volume and FIB-1 abundance. (A) Live confocal images of intestinal nucleoli of day 1 adult worms expressing endogenously-tagged Fibrillarin (wrmScarlet11::FIB-1) and ubiquitous *eft-3p*::wrmScarlet1-10. Scale bar = 5 μm. (B-D) Quantification of the volume (B), concentration (C, mean fluorescence intensity) and total amount (D, total fluorescence intensity) of FIB-1. Each dot represents one nucleolus, and bars and error bars represent mean and 95% confidence interval. P values from Mann-Whitney U tests after Benjamini-Hochberg multiple hypothesis testing correction.

## References

1. Armenti, S.T., L.L. Lohmer, D.R. Sherwood, and J. Nance. 2014. Repurposing an endogenous degradation system for rapid and targeted depletion of C. elegans proteins. Development. 141:4640–4647. doi:10.1242/dev.115048.

2. Berry, S., M. Müller, A. Rai, and L. Pelkmans. 2022. Feedback from nuclear RNA on transcription promotes robust RNA concentration homeostasis in human cells. Cell Systems. 13:454–470.e15. doi:10.1016/j.cels.2022.04.005.

3. Cantwell, H., and P. Nurse. 2019. Unravelling nuclear size control. Curr Genet. doi:10.1007/s00294-019-00999-3.

4. Chen, Y., J.-H. Huang, C. Phong, and J.E. Ferrell. 2023. Protein homeostasis from diffusion-dependent control of protein synthesis and degradation. 2023.04.24.538146. doi:10.1101/2023.04.24.538146.

5. Chen, Y., G. Zhao, J. Zahumensky, S. Honey, and B. Futcher. 2020. Differential Scaling of Gene Expression with Cell Size May Explain Size Control in Budding Yeast. Molecular Cell. 78:359–370.e6. doi:10.1016/j.molcel.2020.03.012.

6. Claude, K.-L., D. Bureik, D. Chatzitheodoridou, P. Adarska, A. Singh, and K.M. Schmoller. 2021. Transcription coordinates histone amounts and genome content. Nat Commun. 12:4202. doi:10.1038/s41467-021-24451-8.

7. Cox, J., and M. Mann. 2012. 1D and 2D annotation enrichment: a statistical method integrating quantitative proteomics with complementary high-throughput data. BMC Bioinformatics. 13:S12. doi:10.1186/1471-2105-13-S16-S12.

8. Darmasaputra, G.S., L.M. van Rijnberk, and M. Galli. 2024. Functional consequences of somatic polyploidy in development. Development. 151:dev202392. doi:10.1242/dev.202392.

9. Dej, K.J., and A.C. Spradling. 1999. The endocycle controls nurse cell polytene chromosome structure during Drosophila oogenesis. Development. 126:293–303. doi:10.1242/dev.126.2.293.

10. Demouchy, F., O. Nicolle, G. Michaux, and A. Pacquelet. 2024. PAR-4/LKB1 prevents intestinal hyperplasia by restricting endoderm specification in Caenorhabditis elegans embryos. Development. 151:dev202205. doi:10.1242/dev.202205.

11. Dickinson, D.J., A.M. Pani, J.K. Heppert, C.D. Higgins, and B. Goldstein. 2015. Streamlined Genome Engineering with a Self-Excising Drug Selection Cassette. Genetics. 200:1035– 1049. doi:10.1534/genetics.115.178335.

12. Edgar, B.A., N. Zielke, and C. Gutierrez. 2014. Endocycles: a recurrent evolutionary innovation for post-mitotic cell growth. Nat Rev Mol Cell Biol. 15:197–210. doi:10.1038/nrm3756.

13. Frank, D.J., and M.B. Roth. 1998. ncl-1 Is Required for the Regulation of Cell Size and Ribosomal RNA Synthesis in Caenorhabditis elegans. Journal of Cell Biology. 140:1321–1329. doi:10.1083/jcb.140.6.1321.

14. Gentric, G., and C. Desdouets. 2014. Polyploidization in Liver Tissue. The American Journal of Pathology. 184:322–331. doi:10.1016/j.ajpath.2013.06.035.

15. Hansson, K.-A., E. Eftestøl, J.C. Bruusgaard, I. Juvkam, A.W. Cramer, A. Malthe-Sørenssen, D.P. Millay, and K. Gundersen. 2020. Myonuclear content regulates cell size with similar scaling properties in mice and humans. Nature Communications. 11:6288. doi:10.1038/s41467-020-20057-8.

16. Hedgecock, E.M., and J.G. White. 1985. Polyploid tissues in the nematode Caenorhabditis elegans. Developmental Biology. 107:128–133. doi:10.1016/0012-1606(85)90381-1.

17. Kaletsky, R., V. Yao, A. Williams, A.M. Runnels, A. Tadych, S. Zhou, O.G. Troyanskaya, and C.T. Murphy. 2018. Transcriptome analysis of adult Caenorhabditis elegans cells reveals tissue-specific gene and isoform expression. PLOS Genetics. 14:e1007559. doi:10.1371/journal.pgen.1007559.

18. Lanz, M.C., E. Zatulovskiy, M.P. Swaffer, L. Zhang, I. Ilerten, S. Zhang, D.S. You, G. Marinov, P. McAlpine, J.E. Elias, and J.M. Skotheim. 2022. Increasing cell size remodels the proteome and promotes senescence. Molecular Cell. 82:3255–3269.e8. doi:10.1016/j.molcel.2022.07.017.

19. Lee, C., H.S. Seidel, T.R. Lynch, E.B. Sorensen, S.L. Crittenden, and J. Kimble. 2017. Single-molecule RNA Fluorescence in situ Hybridization (smFISH) in Caenorhabditis elegans. Bio Protoc. 7:e2357. doi:10.21769/BioProtoc.2357.

20. Lee, L.-W., C.-C. Lee, C.-R. Huang, and S.J. Lo. 2012. The Nucleolus of *Caenorhabditis elegans*. BioMed Research International. 2012:e601274. doi:10.1155/2012/601274.

21. Ma, Y., K. Jonsson, B. Aryal, L. De Veylder, O. Hamant, and R.P. Bhalerao. 2022. Endoreplication mediates cell size control via mechanochemical signaling from cell wall. Science Advances. 8:eabq2047. doi:10.1126/sciadv.abq2047.

22. Marshall, W.F., K.D. Young, M. Swaffer, E. Wood, P. Nurse, A. Kimura, J. Frankel, J. Wallingford, V. Walbot, X. Qu, and A.H. Roeder. 2012. What determines cell size? BMC Biology. 10:101. doi:10.1186/1741-7007-10-101.

23. Mu, L., J.H. Kang, S. Olcum, K.R. Payer, N.L. Calistri, R.J. Kimmerling, S.R. Manalis, and T.P. Miettinen. 2020. Mass measurements during lymphocytic leukemia cell polyploidization decouple cell cycle- and cell size-dependent growth. PNAS. 117:15659–15665. doi:10.1073/pnas.1922197117.

24. Naturale, V.F., M.A. Pickett, and J.L. Feldman. 2023. Persistent cell contacts enable E-cadherin/HMR-1- and PAR-3-based symmetry breaking within a developing *C. elegans* epithelium. Developmental Cell. 58:1830–1846.e12. doi:10.1016/j.devcel.2023.07.008.

25. Neurohr, G.E., R.L. Terry, J. Lengefeld, M. Bonney, G.P. Brittingham, F. Moretto, T.P. Miettinen, L.P. Vaites, L.M. Soares, J.A. Paulo, J.W. Harper, S. Buratowski, S. Manalis, F.J. van Werven, L.J. Holt, and A. Amon. 2019. Excessive Cell Growth Causes Cytoplasm Dilution And Contributes to Senescence. Cell. 176:1083–1097.e18. doi:10.1016/j.cell.2019.01.018.

26. Nonet, M.L. 2020. Efficient Transgenesis in Caenorhabditis elegans Using Flp Recombinase-Mediated Cassette Exchange. Genetics. 215:903–921. doi:10.1534/genetics.120.303388.

27. Nousch, M. 2020. RPL-4 and RPL-9 ̶Mediated Ribosome Purifications Facilitate the Efficient Analysis of Gene Expression in Caenorhabditis elegans Germ Cells. G3 Genes|Genomes|Genetics. 10:4063–4069. doi:10.1534/g3.120.401644.

28. O’Farrell, P.H. 2015. Growing an Embryo from a Single Cell: A Hurdle in Animal Life. Cold Spring Harb Perspect Biol. 7:a019042. doi:10.1101/cshperspect.a019042.

29. Perez, M.F., and B. Lehner. 2019. Vitellogenins - Yolk Gene Function and Regulation in Caenorhabditis elegans. Frontiers in Physiology. 10.

30. Rijnberk, L.M. van, R. Barrull-Mascaró, R.L. van der Palen, E.S. Schild, H.C. Korswagen, and M. Galli. 2022. Endomitosis controls tissue-specific gene expression during development. PLOS Biology. 20:e3001597. doi:10.1371/journal.pbio.3001597.

31. Roeder, A.H.K., A. Cunha, C.K. Ohno, and E.M. Meyerowitz. 2012. Cell cycle regulates cell type in the Arabidopsis sepal. Development. 139:4416–4427. doi:10.1242/dev.082925.

32. Sallee, M.D., M.A. Pickett, and J.L. Feldman. 2021. Apical PAR complex proteins protect against programmed epithelial assaults to create a continuous and functional intestinal lumen. eLife. 10:e64437. doi:10.7554/eLife.64437.

33. Sallee, M.D., J.C. Zonka, T.D. Skokan, B.C. Raftrey, and J.L. Feldman. 2018. Tissue-specific degradation of essential centrosome components reveals distinct microtubule populations at microtubule organizing centers. PLoS Biol. 16. doi:10.1371/journal.pbio.2005189.

34. Schmoller, K.M., J.J. Turner, M. Kõivomägi, and J.M. Skotheim. 2015. Dilution of the cell cycle inhibitor Whi5 controls budding-yeast cell size. Nature. 526:268–272. doi:10.1038/nature14908.

35. Sen, I., A. Kavšek, and C.G. Riedel. 2021. Chromatin Immunoprecipitation and Sequencing (ChIP-seq) Optimized for Application in Caenorhabditis elegans. Current Protocols. 1:e187. doi:10.1002/cpz1.187.

36. Swaffer, M.P., J. Kim, D. Chandler-Brown, M. Langhinrichs, G.K. Marinov, W.J. Greenleaf, A. Kundaje, K.M. Schmoller, and J.M. Skotheim. 2021. Transcriptional and chromatin-based partitioning mechanisms uncouple protein scaling from cell size. Molecular Cell. 81:4861–4875.e7. doi:10.1016/j.molcel.2021.10.007.

37. Swaffer, M.P., G.K. Marinov, H. Zheng, L. Fuentes Valenzuela, C.Y. Tsui, A.W. Jones, J. Greenwood, A. Kundaje, W.J. Greenleaf, R. Reyes-Lamothe, and J.M. Skotheim. 2023. RNA polymerase II dynamics and mRNA stability feedback scale mRNA amounts with cell size. Cell. 186:5254–5268.e26. doi:10.1016/j.cell.2023.10.012.

38. Unhavaithaya, Y., and T.L. Orr-Weaver. 2012. Polyploidization of glia in neural development links tissue growth to blood–brain barrier integrity. Genes Dev. 26:31–36. doi:10.1101/gad.177436.111.

39. Uppaluri, S., S.C. Weber, and C.P. Brangwynne. 2016. Hierarchical Size Scaling during Multicellular Growth and Development. Cell Reports. 17:345–352. doi:10.1016/j.celrep.2016.09.007.

40. Van Rompay, L., C. Borghgraef, I. Beets, J. Caers, and L. Temmerman. 2015. New genetic regulators question relevance of abundant yolk protein production in C. elegans. Sci Rep. 5:16381. doi:10.1038/srep16381.

41. Weigert, M., U. Schmidt, R. Haase, K. Sugawara, and G. Myers. 2020. Star-convex Polyhedra for 3D Object Detection and Segmentation in Microscopy. *In* 2020 IEEE Winter Conference on Applications of Computer Vision (WACV). 3655–3662.

42. Xie, S., and J.M. Skotheim. 2020. A G1 Sizer Coordinates Growth and Division in the Mouse Epidermis. Current Biology. 30:916–924.e2. doi:10.1016/j.cub.2019.12.062.

43. Xiong, R., and K. Sugioka. 2020. Improved 3D cellular morphometry of Caenorhabditis elegans embryos using a refractive index matching medium. PLOS ONE. 15:e0238955. doi:10.1371/journal.pone.0238955.

44. Yi, Y.-H., T.-H. Ma, L.-W. Lee, P.-T. Chiou, P.-H. Chen, C.-M. Lee, Y.-D. Chu, H. Yu, K.-C. Hsiung, Y.-T. Tsai, C.-C. Lee, Y.-S. Chang, S.-P. Chan, B.C.-M. Tan, and S.J. Lo. 2015. A Genetic Cascade of let-7-ncl-1-fib-1 Modulates Nucleolar Size and rRNA Pool in Caenorhabditis elegans. PLOS Genetics. 11:e1005580. doi:10.1371/journal.pgen.1005580.

45. Zeldovich, V.B., C.H. Clausen, E. Bradford, D.A. Fletcher, E. Maltepe, J.R. Robbins, and A.I. Bakardjiev. 2013. Placental Syncytium Forms a Biophysical Barrier against Pathogen Invasion. PLOS Pathogens. 9:e1003821. doi:10.1371/journal.ppat.1003821.

